# Image-based Phenotyping and Genetic Analysis of Potato Skin Set and Color

**DOI:** 10.1101/694745

**Authors:** Maria V. Caraza-Harter, Jeffrey B. Endelman

## Abstract

Image-based phenotyping offers new opportunities for fast, objective, and reliable measurement for breeding and genetics research. In the current study, image analysis was used to quantify potato skin color and skin set, which are critical for the marketability of new varieties. A set of 15 red potato varieties and advanced breeding lines was evaluated over two years at a single location, with two harvest times in the second year. After mechanical harvest and grading, 7-8 representative tubers per plot were photographed, and the photos were analyzed with ImageJ to measure skinning (as % surface area) and skin color using the Hue, Chroma and Lightness (HCL) representation. The plot-based heritability was consistently high (> 0.77) across traits and environments; the genetic correlation between environments was also high, ranging from 0.81 to 0.98. Significant increases in Lightness and Chroma, as well as a decrease in skinning, were observed at the late compared to early harvest, while the opposite trends for color were observed after six weeks of storage. The three color traits were unexpectedly collinear in this study, with the first principal component explaining 86% of the variation. This result may reflect the physiology of red color in potato, but the highly selected nature of the 15 genotypes may also be a factor. Image-based phenotyping offers new opportunities to advance genetic gain and understanding for tuber appearance traits that have been difficult to precisely measure in the past.

---

Potato is an important crop for both processing and fresh markets, with the latter category representing 27% of total US production. The main fresh market types sold in the US are russets, whites, yellows, and reds. Tuber appearance has a strong impact on marketability and is therefore important to evaluate during variety development. Historically, potato breeders have used visual ratings (i.e., 1–5) to score traits that affect tuber appearance, such as length/width ratio, height/width ratio, curvature, eye depth, skin color, skin finish (i.e., netted vs. smooth), and skin set (i.e., resistance to excoriation). This approach is labor-intensive, subjective, and often lacks precision. The alternative of image-based phenotyping of tuber appearance provides an opportunity to move beyond these limitations. Computer vision has been previously used in potato and other horticultural crops for grading of produce, such as identifying defects in shape or color (Patel et al., 2012; Tao et al., 1995). Instead of quality control purposes, our motivation is to investigate the genetics of skin set and color in red potatoes.

Red skin color in potatoes is due to the presence of anthocyanin pigments in the tuber periderm, which can be quantified to study the influence of variety and management on color (Andersen et al., 2002; Hung et al., 1997; Roe et al., 2014; Rosen et al., 2009; Waterer, 2010). With image-based phenotyping, however, the goal is to measure human perception of color rather than its chemical basis. A number of mathematical models exist for representing color. The RGB (red, green, blue) model is widely known and used in digital cameras, but the biconic Hue, Chroma and Lightness (HCL) model is more closely related to human perception (Figure 1). Hue corresponds to the polar angle, which we have centered on red at 0°, with yellow at 60° and magenta at -60°. The vertical dimension is Lightness, ranging from 0 (black) to 1 (white). Chroma is the radial dimension, which ranges from grayscale at 0 to fully saturated at C = 2L. The HCL color model has been used before in potato based on measurements with a handheld colorimeter, to study the effects of management, soil type, and storage on a limited number of varieties (Andersen et al., 2002; Roe et al., 2014; Rosen et al., 2009). The current study builds on this earlier research by (1) extracting HCL phenotypes from images and (2) examining a larger set of genotypes.

**Figure 1.**
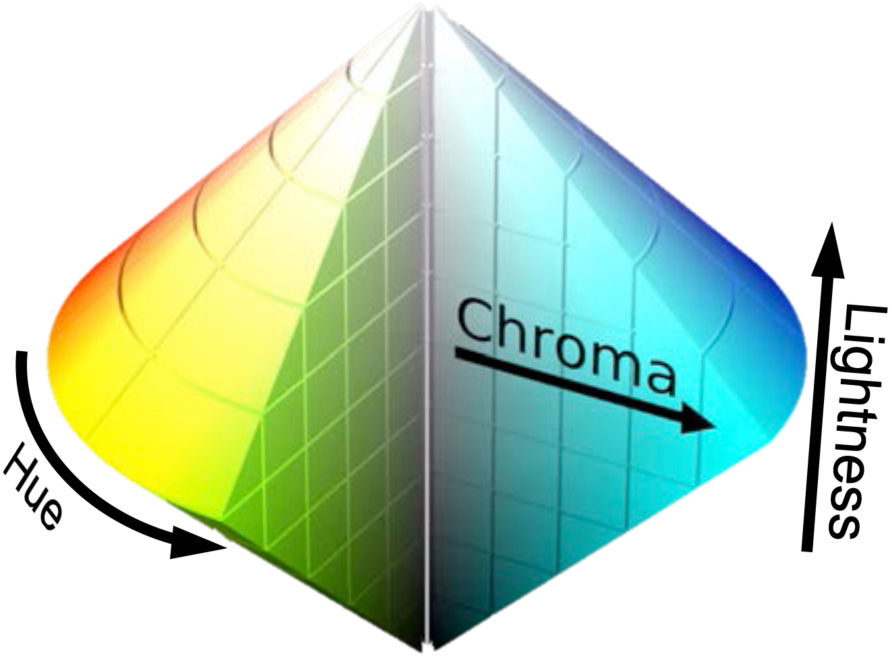
Biconic geometry of the HCL color model. Image distributed under CC BY-SA 3.0 (Jacob Rus, SharkD, https://commons.wikimedia.org/wiki/File:HSL_color_solid_dblcone_chroma_gray.png; changed background to white and text angle).

Potato tuber “skin,” or periderm, is composed of three tissues: phellem, phellogen and phelloderm (Reeve et al., 1969). The phellogen is meristematic (cork cambium) tissue, adding cells of suberized phellem to the outside and phelloderm tissue to the inside (Lulai et al., 2001). During the early stages of tuber development, the phellogen layer is active and very susceptible to excoriation, or “skinning.” Skinning not only reduces the marketability of tubers but also the ability to retain moisture and resist disease during storage. As the tuber matures, changes in the phellogen layer promote greater adhesion of the phellem—a process informally known as skin set. According to USDA grading standards, the highest grade of “practically no skinning” means not more than 5% of the potatoes have more than 10% of the skin missing or feathered (USDA, 2008).

Two different approaches to measuring skin set have been used. One is to measure the torque at which the periderm excoriates using a torquemeter (Lulai et al., 1993). A more direct approach, which is amenable to image-based phenotyping and more closely aligned with human perception, is to record the percent of missing skin on an area basis (Gao et al., 2016). Varietal differences in skin set are widely recognized, but very little is known about the genetic basis of this critical trait (Halderson et al., 1993; Lulai, 2007).

The objectives of this study were to (1) use image-based phenotyping to measure skin color and skin set for a group of 15 commercial varieties and advanced breeding lines; (2) determine the heritability of the image-based phenotypes, both within and across environments; and (3) investigate the influence of harvest and storage time on these traits.

## MATERIALS AND METHODS

### Plant Material and Field Trials

A group of 15 red varieties and advanced breeding lines from the University of Wisconsin-Madison were evaluated in 2015 and 2016 as replicated 15-plant plots at the UW Hancock Agricultural Research Station. We evaluated 13 clones in 2015 using a randomized complete block design (RCBD) with three replications. The experiment was planted April 27 and harvested 121 Days After Planting. Fertility, water, and pest management followed UW-Extension guidelines for potato (Bussan et al., 2015). Diquat bromide was applied 14 and 7 days before harvest to promote vine desiccation. Tubers were mechanically harvested into 30 cm × 45 cm rigid plastic milk crates, run through a washing and grading line, and then crated up again for storage at 12°C with 95% relative humidity. No additional steps were taken to promote skinning. In 2016, 11 of the 13 clones from the 2015 trial were evaluated again, plus two additional check varieties, for a total of 13 clones (Table S1). The 2016 experiment consisted of two adjacent RCBD trials, each with two replicates, planted on April 21. The first trial was harvested 109 DAP and the second 138 DAP. Crop management and harvest followed the same protocols as 2015.

### Image Acquisition and Analysis

Photos were taken within a few days of harvest in 2015 and 2016, as well as six weeks after harvest in 2016. A set of 7-8 representative tubers from each plot were placed on a black board and photographed, on one side in 2015 and on both sides in 2016, using a Photosimile 200 Lightbox equipped with a CanonEOS T5i camera (Figure S1). The camera was set to autofocus with an aperture of F20, an ISO 100 and a shutter speed of 1/10. A Small MacBeth Color Card was included in each photo in 2016 to compensate for potential variation in lighting and exposure.

The dataset of 253 images was analyzed using the ImageJ software (Schneider et al., 2012). For the 2016 photos, the first step consisted of image calibration based on the color card, using the ImageJ plugin Chart White Balance (Vander Haeghen, 2007). For both years, a semi-automated background removal was performed using color thresholds to differentiate tubers from the black background, and the total tuber surface area was measured in pixels. Hue and brightness thresholds were then used to select and measure the skinned surface area. The ratio between the skinned area and total tuber surface area is reported as skinning percentage (%) to quantify skin set. To measure color, hue and brightness thresholds were used to select red skin and exclude external defects such as exposed tissue (due to skinning) and common scab. The RGB Measure Plugin was used to measure the average R, G, and B values of the selected area on a 0–255 scale. RGB values were divided by 255 to fall in the range 0–1 and then converted to the HCL representation according to the following standard formulas (Smith, 1978):

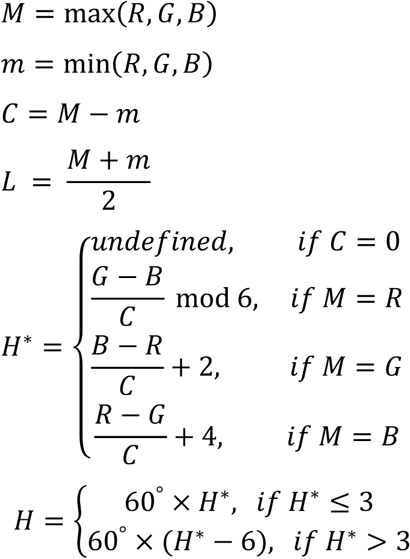

### Statistical Analysis

Initially, the color and skin set measurements taken at harvest were analyzed separately for each of the three environments: 2015@121DAP, 2016@109DAP, and 2016@138DAP. The phenotype *P*_*ij*_ for genotype *i* in block *j* was modeled by

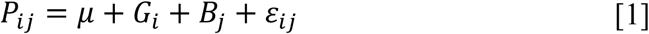

where *μ* is the intercept and *G*_*i*_, *B*_*j*_, and *ε*_*ij*_ are normally distributed random effects for genotype, block and residuals, respectively. Variance components were estimated by Restricted Maximum Likelihood with the ASReml-R software (Butler et al., 2009; R. Core Team, 2018). After inspecting the residuals, a log transformation was used for skinning % to satisfy the normality assumption. Plot-based heritability was estimated by

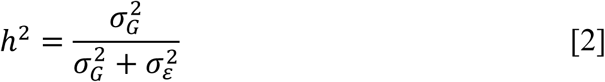

where 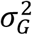 and 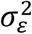 are the variance components for genotype and residual, respectively.

For the combined analysis of the at-harvest measurements from the three environments, we fitted the following mixed model:

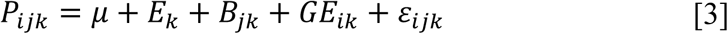

In Equation 3, *μ* is the intercept, *E*_*k*_ is the fixed effect of environment, *B*_*jk*_ is the random effect of block nested within environment, *GE*_*ik*_ is the random effect of genotype *i* nested within environment *k*, and *ε*_*ijk*_ are residuals. A separable covariance model was used for the *GE*_*ik*_ effect, such that the effects for two different genotypes were independent but not the effects for the same genotype in two different environments:

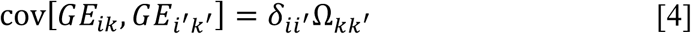

In Equation 4, *δ*_*ii*_′ is the Kronecker delta, which equals 1 when its two arguments are identical and 0 otherwise, and Ω_*kk*_′ is the genetic covariance between environments *k* and *k’*. Variance components and the fixed effects for environment were estimated using ASReml-R. The statistical significance of pairwise differences between environments was determined with ASReml-R based on a Wald test and *p* = 0.05 threshold. The genetic correlation ρ_*kk*_′ between environments *k* and *k’* was calculated as

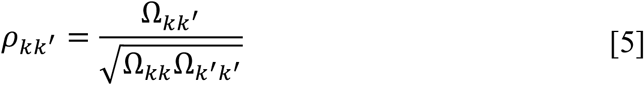

Because of the high genetic correlation between environments, a single BLUP (best unbiased linear predictor) was calculated for each clone using a modification of Equation 3. The *GE*_*iK*_ effect was rewritten as 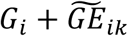 to separate the main effect from the G×E interaction, both of which were assumed to be normally distributed and independent: 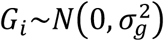 and 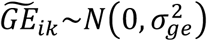. Variance components were estimated with ASReml-R and BLUP[*G*_*i*_] ≡ *Ĝ*_*i*_ was calculated from the Henderson (1975) mixed model equations. The reliability 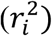 of *Ĝ*_*i*_ was estimated from its prediction error variance (*PEV*_*i*_ = *Var*[*Ĝ*_*i*_ − *G*_*i*_]) according to (Clark et al., 2012)

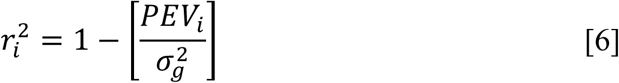

Principal component analysis of the three color trait BLUPs, standardized to have zero mean and unit variance, was performed using the *princomp* function in R.

The effect of storage time on the color traits was estimated using the 2016 data, based on the following linear mixed model:

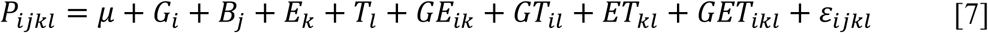

In Equation 7, the intercept is represented by *μ*; *G*_*i*_, *B*_*j*_, and ε_*ijk*_ are the random effects for genotype, block, and residuals respectively. *E*_*k*_ is the fixed effect of environment, with two levels for the factor (109 and 138 DAP), and *T*_*l*_ is the fixed effect of storage time, with two levels for the factor (0 and 6 weeks after harvest). All interaction terms were random except *ET*_*kl*_. Because the effect of storage time was estimated from measurements on the same field plot, a correlated model for the residuals was used:

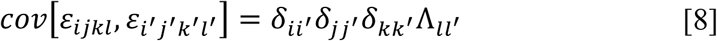

In Equation 8, *δ* is the Kronecker delta, and Λ is a 2×2 covariance matrix estimated with ASReml-R. The statistical significance of the storage time effect (*T*_*l*_) was determined based on a Wald test and *p* = 0.05 threshold. From Equation 7, the intraclass correlation ρ_*ll*_′ between the genotypic values of one clone at different storage times (from the same field environment) is

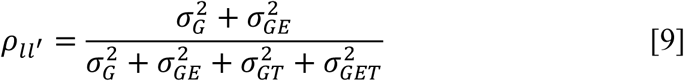

## RESULTS

In the first experiment (in 2015), photographs of 13 red clones were taken within a few days of harvest (121 DAP) and used to estimate Hue, Chroma, Lightness, and skinning % for each plot (Figure 2). Hue ranged from -4.6° to 6.5°, Chroma from 0.26 to 0.35, and Lightness from 0.25 to 0.33. The range for skinning was 2.2% to 30.2%. The plot-based heritability exceeded 0.75 for all four traits (Table 1). As shown in Figure 2, the three color traits were highly correlated, but skinning showed only weak or no correlation with color.

**Table 1.**
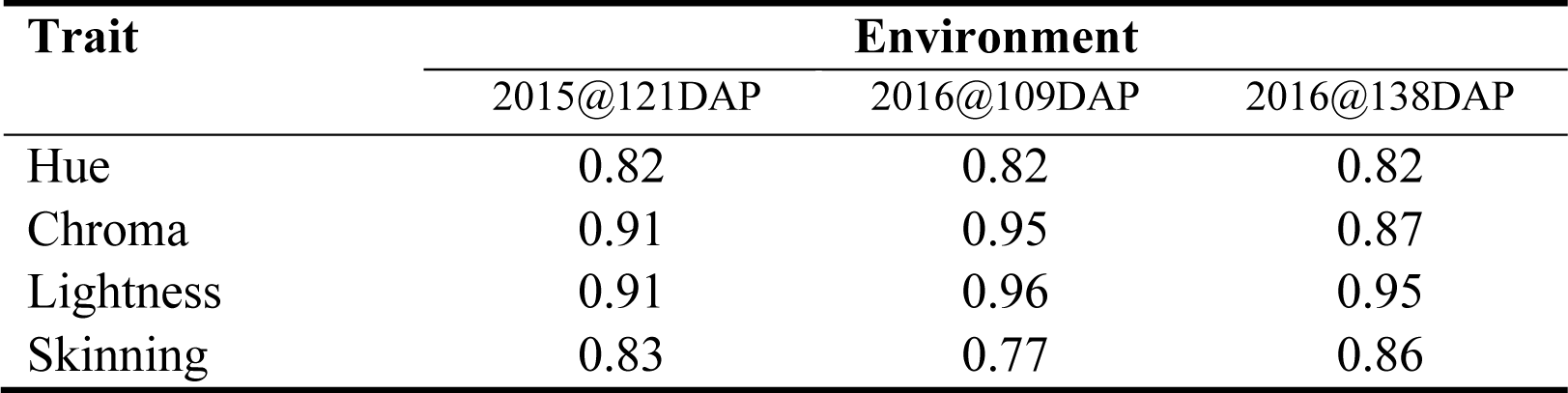
Plot-based heritability for color and skin set measurements taken at harvest, for three environments (Year@HarvestTime) in Wisconsin.

**Figure 2.**
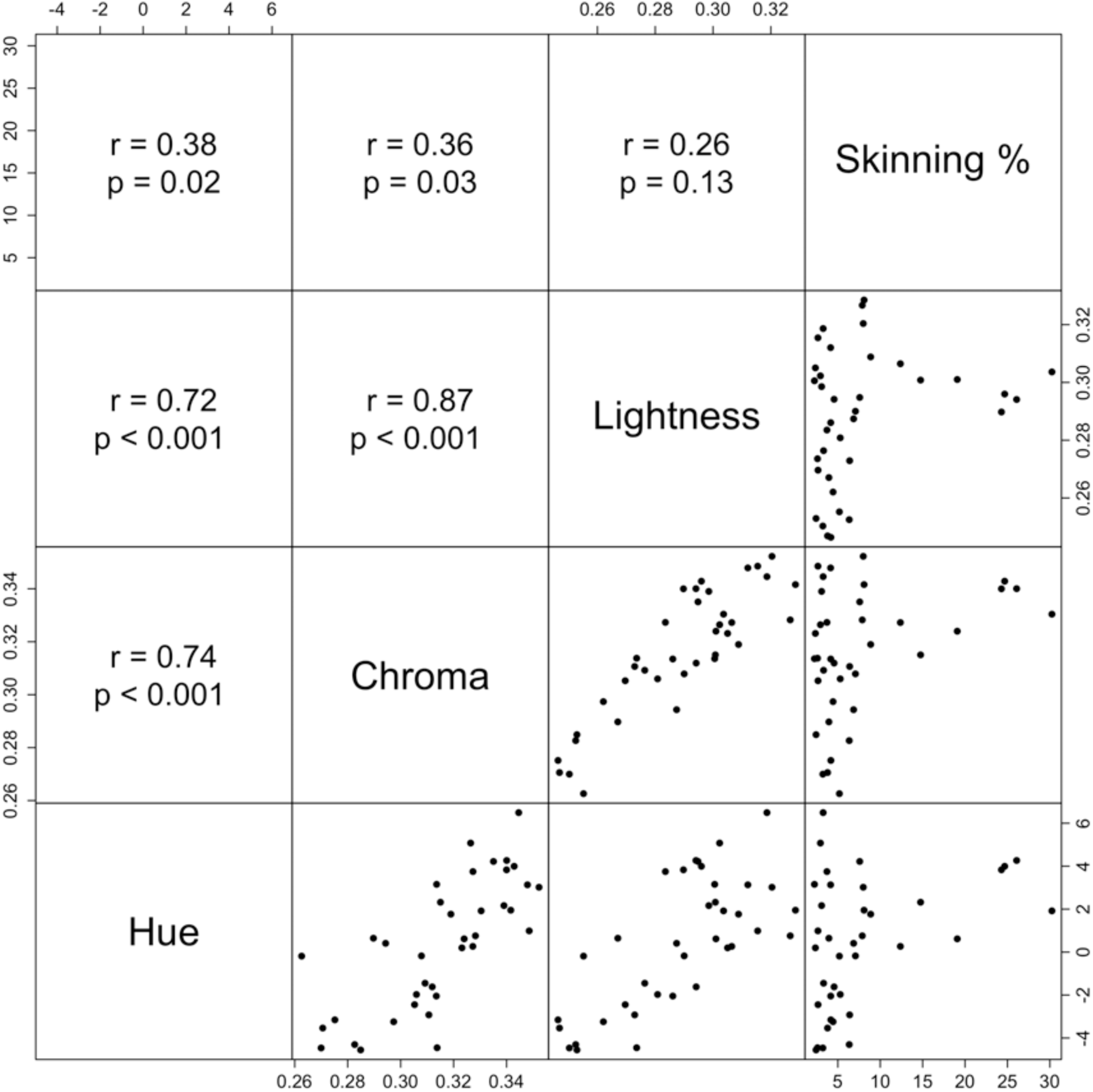
Pairwise scatterplot of Hue, Chroma, Lightness and Skinning % for each plot of the 2015@121DAP experiment. Correlation coefficient (*r*) and p-values shown above the diagonal.

The experiment was repeated for a second season, in 2016, with separate trials for early (109 DAP) and late (138 DAP) harvests. The plot-based heritability for both harvest times was similar to the 2015 experiment (Table 1). From a combined analysis of the three environments (2015@121DAP, 2016@109DAP, 2016@138DAP), the statistical significance of the environment effect was estimated (Table 2). Hue was significantly higher and Lightness significantly lower in 2015 compared to 2016. Looking at the effect of harvest time in 2016, there was no significant difference in Hue, while Chroma and Lightness were both higher for the late harvest. Skinning was not significantly different between 2015 and the early 2016 harvest, but less skinning was observed in the late 2016 harvest.

**Table 2.** Environment means for the color traits and skin set. Means with different letters are significantly different based on a Wald test with *p* < 0.05.

| Trait | Environment |  |  |
| --- | --- | --- | --- |
|  | 2015@121DAP | 2016@109DAP | 2016@138DAP |
| Hue (°) | 1.02<sup>a</sup> | -2.57<sup>b</sup> | -2.35<sup>b</sup> |
| Chroma | 0.32<sup>a</sup> | 0.32<sup>a</sup> | 0.34<sup>b</sup> |
| Lightness | 0.29<sup>a</sup> | 0.33<sup>b</sup> | 0.35<sup>c</sup> |
| Skinning (%) | 5.89<sup>a</sup> | 4.99<sup>a</sup> | 3.26<sup>b</sup> |

Despite the significant main effect of environment, there was very little G×E in this experiment. The genetic correlation between environments exceeded 0.8 for all four traits and all three pairwise comparisons (Table 3). This allowed for the calculation of a single BLUP per clone across the three environments (Table S2). For the clones evaluated in both years, the reliability of the BLUPs exceeded 0.7 for all four traits (Table S2). Because of the high correlation between the three color traits (Figure 1), a principal component (PC) analysis of the BLUPs was performed. The first PC captured 83% of the variation (Figure S2), with loadings of 0.57 for Hue and Chroma and 0.60 for Lightness.

**Table 3.** Genetic correlations for color and skin set between three Wisconsin environments, based on measurements taken at harvest.

| Trait | 2015@121DAP–<br>2016@109DAP | 2015@121DAP–<br>2016@138DAP | 2016@109DAP–<br>2016@138DAP |
| --- | --- | --- | --- |
| Hue | 0.81 | 0.83 | 0.98 |
| Chroma | 0.87 | 0.82 | 0.88 |
| Lightness | 0.85 | 0.93 | 0.96 |
| Skinning | 0.89 | 0.96 | 0.84 |

The first PC for color was plotted against skinning % to visualize the genetic variation for this set of 15 clones (Figure 3). The top two red varieties in Wisconsin, as well as the entire US, are Red Norland and Dark Red Norland (National Potato Council, 2018), which are line selections (i.e., somatic mutants) of Norland (Johansen et al., 1959). A major reason for the continued dominance of Norland selections is resistance to skinning, which is consistent with their position along the horizontal axis in Figure 3. As the name suggests, Dark Red Norland was darker than Red Norland in our experiment, but several breeding lines (e.g., W8893-1R, W6511-1R) were even darker. The potential for large differences even among close relatives is illustrated by the full-sibs W10209-2R and W10209-7R, for which the skinning percentages were 3.2% and 17.1%, respectively.

**Figure 3.**
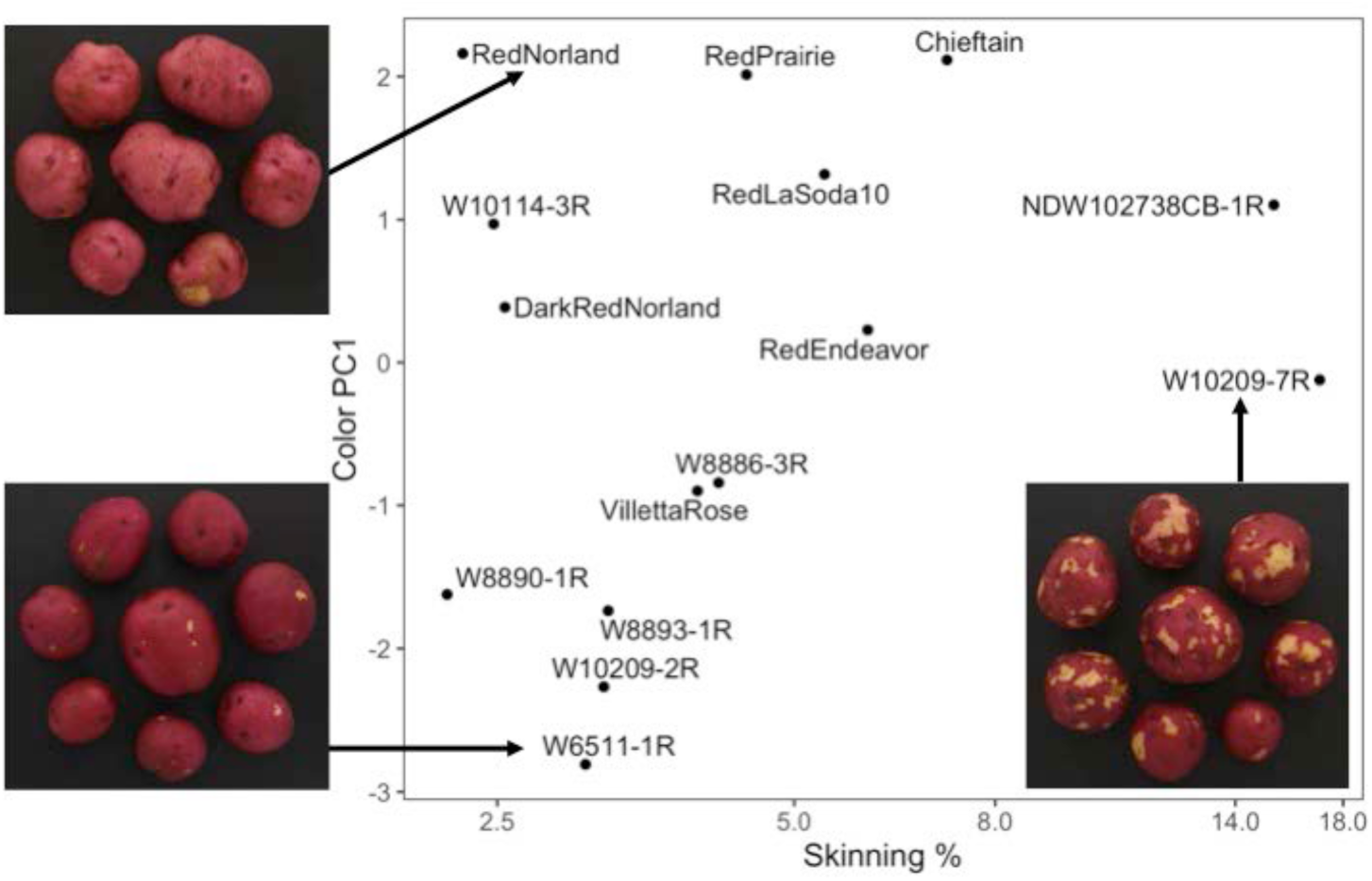
Scatterplot of BLUPs for color (PC1) vs. skinning % for the 15 red clones evaluated in three environments.

The effect of storage time on color was estimated using the 2016 data, for which photographs were taken at 0 and 6 weeks after harvest. For all three color traits, the plot-based heritability remained high after six weeks of storage (≥ 0.88). There was also very little G×E between the two storage times, with genetic correlations above 0.9 for all three traits (Table S3). The color traits were significantly affected by storage time: Hue increased by 3°, Chroma decreased by 0.03, and Lightness decreased by 0.03 (Table 4). The perceived effect of harvest and storage time is visible in Figure 4, which compares images of the variety ‘Red Prairie’ at different harvest times and before and after storage. The BLUPs for each clone at each harvest and storage time are provided in Table S4.

**Table 4.** The effect of storage time on color, from the 2016 experiment. Means with different letters are significantly different based on a Wald test with *p* < 0.05.

| Trait | At Harvest | 6 Weeks |
| --- | --- | --- |
| Hue (°) | -2.49<sup>a</sup> | 0.77<sup>b</sup> |
| Chroma | 0.33<sup>a</sup> | 0.30<sup>b</sup> |
| Lightness | 0.34<sup>a</sup> | 0.31<sup>b</sup> |

**Figure 4.**
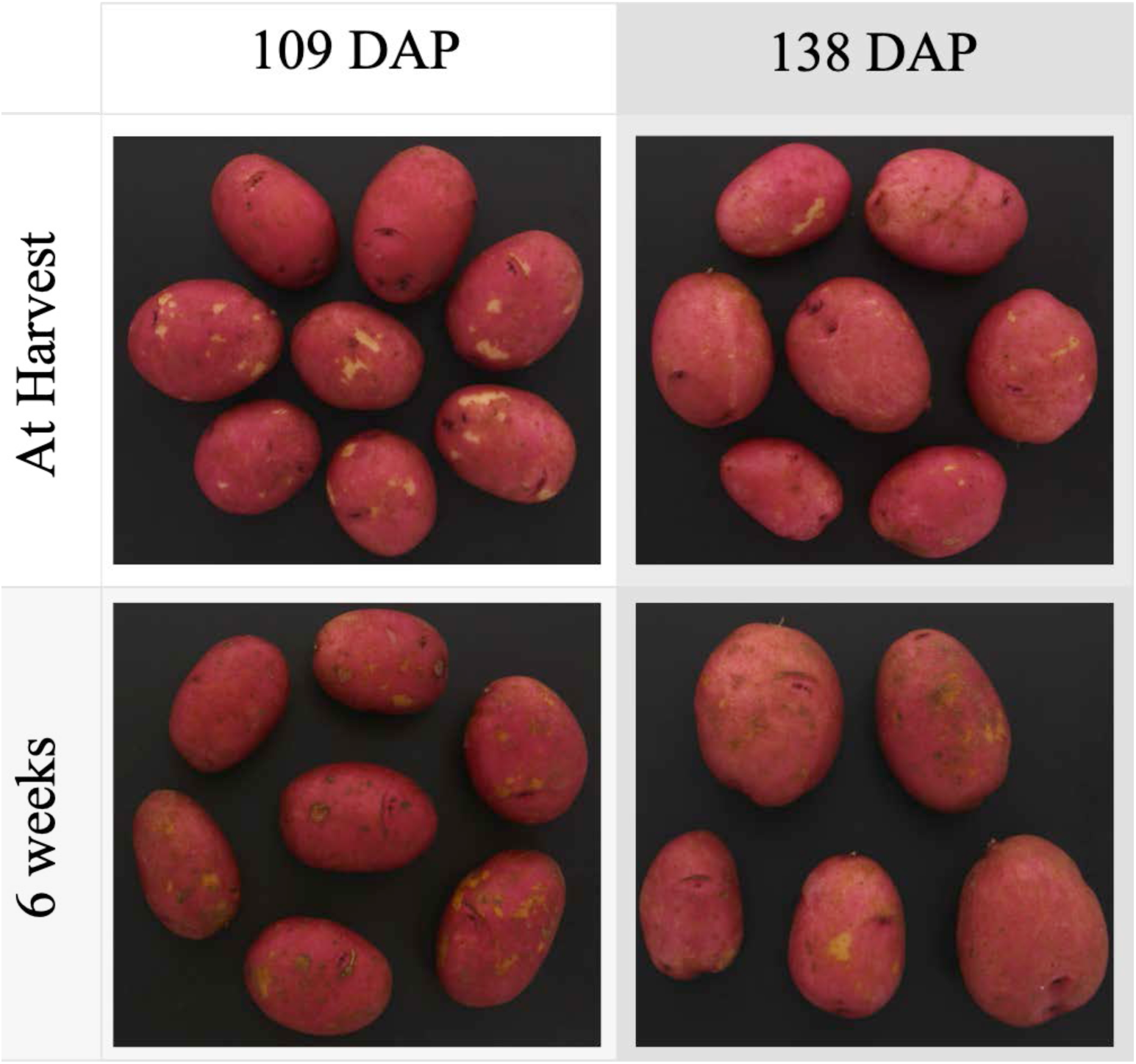
The effects of harvest time (109 vs 138 DAP) and storage time (0 vs. 6 weeks) for the variety ‘Red Prairie.’ Storage led to decreased Chroma and Lightness, while the opposite trends were observed for the late vs. early harvest. Besides the effect of storage on skin color, the images show changes in the color of the skinned area and higher severity of skin blemish diseases (e.g., silver scurf and black dot).

## DISCUSSION

The primary motivation for this research was to develop an image-based phenotyping method for tuber appearance that can be used for breeding and genetics research. Image-based phenotyping is ubiquitous now due to the availability of multi-spectral sensors on UAVs (Li et al., 2019), but imaging studies of plant morphology are also becoming more common (Darrigues et al., 2008; Miller et al., 2017; Moore et al., 2013). Plot-based heritability (*h*^2^) is a critical measure of the reliability of the phenotyping method, and we were pleased to estimate values over 0.75 for all three color traits and skinning percentage. Because *h*^2^ was similar between the early and late harvest in 2016, and because there was very little G×E between these environments (genetic correlations exceeded 0.8), it appears the timing of harvest (within reason) is not critical for selection or genetic mapping for these traits.

Potato growers often refer to the “loss of color” that occurs during storage, which negatively impacts the marketability of red potatoes. In this experiment, “loss of color” manifested as lower Chroma and lower Lightness at 6 weeks after harvest compared to right after harvest (see Table 4). Figure 4 illustrates these changes in tuber appearance for one genotype. Previous studies on the effect of storage on red color, based on measurements with a handheld colorimeter, have also reported decreases in Chroma and/or Lightness (Andersen et al., 2002; Roe et al., 2014; Rosen et al., 2009). Selecting genotypes that maintain Chroma in storage is an important breeding goal, but there was very little G×E for this trait (genetic correlations exceeded 0.9). Since the 15 clones in this experiment are representative of the genetic diversity of the UW-Madison red breeding program, new germplasm may be needed to make genetic gains for color retention.

Compared with storage time, studies on the effect of harvest time on red skin color are rarer and less consistent. In this study, Lightness and Chroma were significantly higher for tubers harvested 138 DAP compared to 109 DAP. Rosen et al. (2009) measured skin color at harvest compared to vine kill in two years, reporting decreased Lightness and no change in Chroma in the first year but higher Lightness and lower Chroma in the second year.

The physiological basis for the changes in tuber appearance reported here deserves further study. Hung et al. (1997) reported that both Chroma and anthocyanidin (the aglycone form of anthocyanin) content per unit surface area decreased during tuber growth (i.e., “bulking”); the authors hypothesized this was due to pigment dilution (from increased surface area) and/or degradation. Sulc et al. (2017) measured anthocyanidin content in potatoes with pigmented skin and flesh (which are a specialty item, not a major commodity, in the US), reporting a fairly consistent decline over a 15-week period. Extrapolating these results to our study, we would predict there to be less anthocyanin in the late-harvest tubers compared to early-harvest, and yet the late-harvest tubers had higher Chroma. Both Andersen et al. (2002) and Roe et al. (2014) reported decreases in anthocyanin content during storage, which seems consistent with our finding of lower Chroma.

The strong collinearity between the three color traits in this study was unexpected. The HCL color model is three-dimensional, but for the 15 genotypes in this study, the color variation was largely one-dimensional (the first PC explained 86% of the variation). This result may reflect the biology of red color in potato, but the highly selected nature of the 15 genotypes in this study may be a factor. Support for the latter hypothesis comes from an ongoing genetic mapping project in which hundreds of unselected F1 progeny from the UW-Madison red potato breeding program have been imaged, and for which the color traits are less correlated (data not shown). The genetics of red skin color as a qualitative (presence/absence) trait is well characterized (Jung et al., 2009; Zhang et al., 2009), but our understanding of color as a quantitative trait, particularly in tetraploid potato, is incomplete. Much less is known about the genetics of skin set, as there have been only a few studies based on gene expression (Neubauer et al., 2013; Vulavala et al., 2017) and none based on association or linkage analysis.

## ACKNOWLEDGMENTS

Financial support was provided by the Wisconsin Department of Agriculture Specialty Crop Block Grant (16-02), the Wisconsin Potato and Vegetable Growers Association, and the UW Office of the Vice Chancellor for Research and Graduate Education. L. Snodgrass, G. Christensen, and B. Kleven assisted with the harvest and photographing of tubers.

## SUPPLEMENTAL MATERIAL

Image-based Phenotyping and Genetic Analysis of Potato Skin Set and Color Caraza-Harter and Endelman

**Figure S1.**
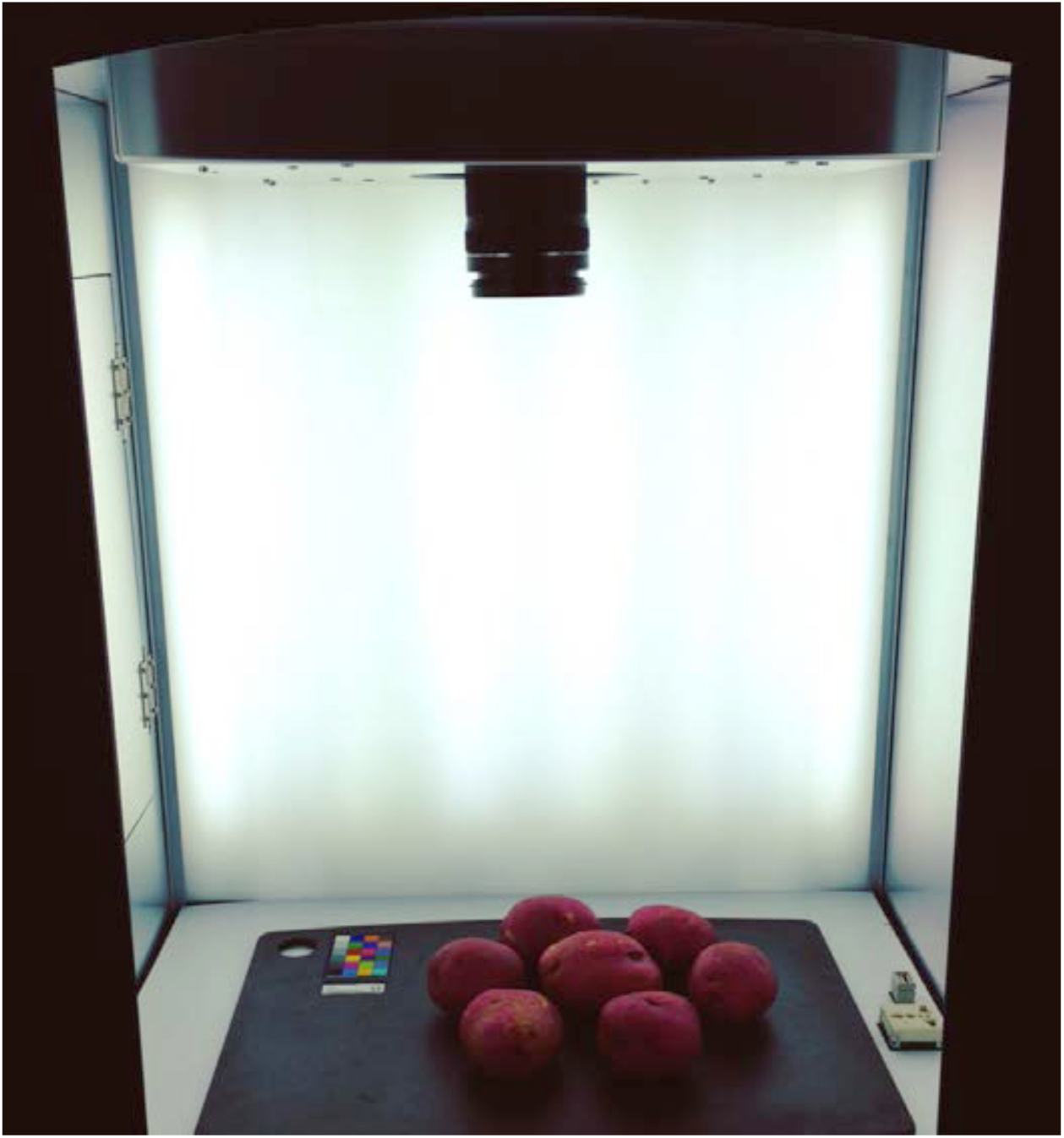
Imaging system at the Hancock Agricultural Research Station in Wisconsin.

**Figure S2.**
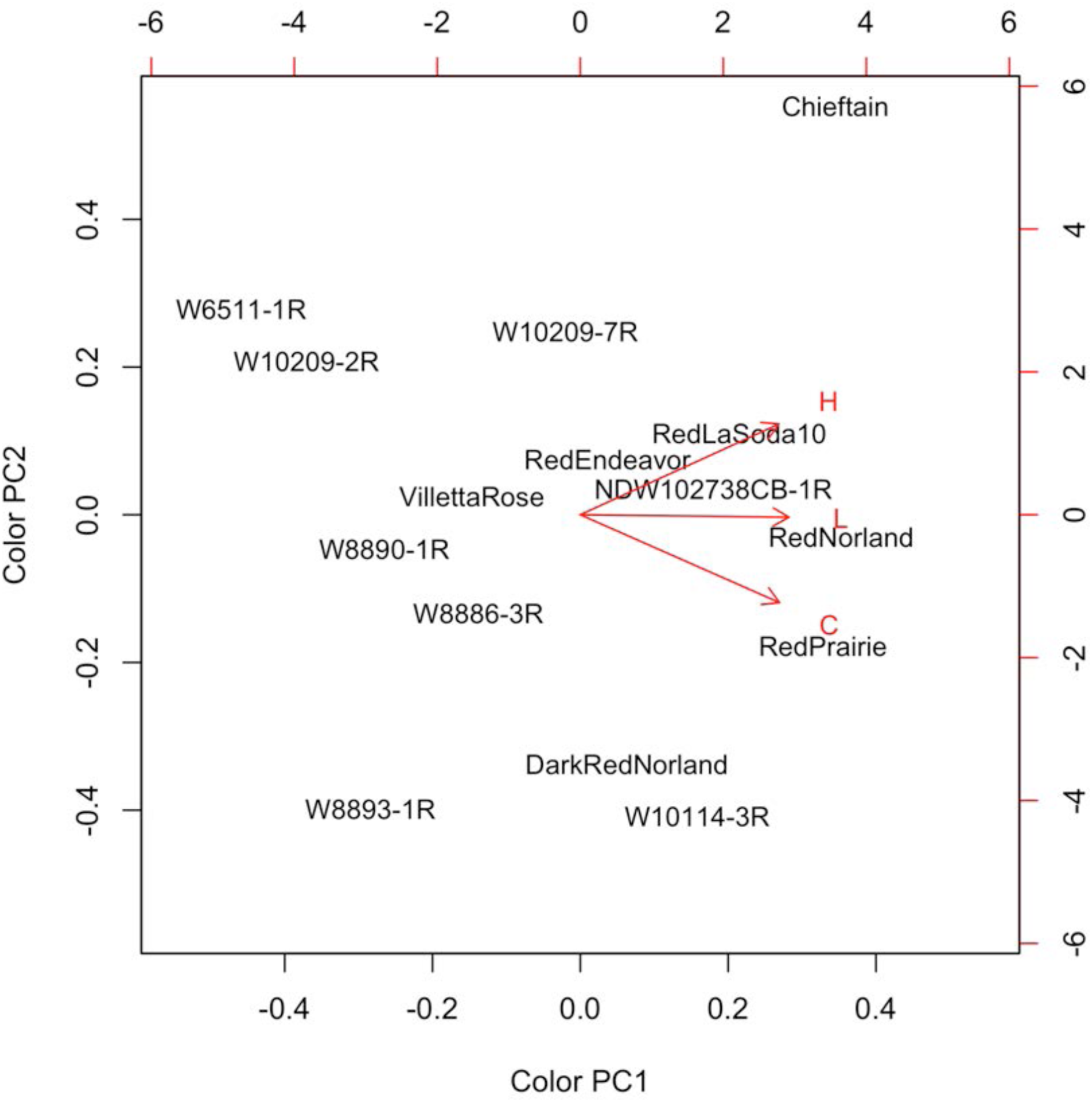
Biplot from a principal component analysis of the BLUPs for the color traits, across all three environments.

**Table S1.** List of red potato varieties and advanced breeding lines evaluated in 2015 and 2016 at the Hancock Agricultural Research Station in Wisconsin.

| Clone | Environment |  |  |
| --- | --- | --- | --- |
|  | 2015 | 2016 |  |
|  | 121 DAP | 109 DAP | 138 DAP |
| Chieftain | NA | * | * |
| DarkRedNorland | * | * | * |
| NDW102738CB-1R | * | NA | NA |
| RedEndeavor | * | * | * |
| RedLaSoda10 | * | * | * |
| RedNorland | NA | * | * |
| RedPrairie | * | * | * |
| VillettaRose | * | * | * |
| W10114-3R | * | * | * |
| W10209-2R | * | * | * |
| W10209-7R | * | * | * |
| W6511-1R | * | * | * |
| W8886-3R | * | NA | NA |
| W8890-1R | * | * | * |
| W8893-1R | * | * | * |

\* : Evaluated
NA: Not Available

**Table S2.** BLUPs and reliabilities across all three environments for the two-year experiment.

| Clone | Hue (°) |  | Chroma |  | Lightness |  | Skinning (%) |  |
| --- | --- | --- | --- | --- | --- | --- | --- | --- |
| | BLUP | $r^2$ | BLUP | $r^2$ | BLUP | $r^2$ | BLUP | $r^2$ |
| Chieftain | 3.35 | 0.77 | 0.33 | 0.84 | 0.35 | 0.88 | 7.15 | 0.82 |
| DarkRedNorland | -2.16 | 0.82 | 0.34 | 0.87 | 0.33 | 0.90 | 2.55 | 0.86 |
| NDW102738CB-1R | 0.97 | 0.68 | 0.35 | 0.77 | 0.32 | 0.84 | 15.31 | 0.76 |
| RedEndeavor | -1.01 | 0.82 | 0.32 | 0.87 | 0.33 | 0.90 | 5.94 | 0.86 |
| RedLaSoda10 | 0.88 | 0.82 | 0.34 | 0.87 | 0.34 | 0.90 | 5.37 | 0.86 |
| RedNorland | 0.38 | 0.78 | 0.34 | 0.84 | 0.38 | 0.88 | 2.31 | 0.83 |
| RedPrairie | 0.39 | 0.82 | 0.35 | 0.87 | 0.36 | 0.90 | 4.47 | 0.86 |
| VillettaRose | -2.67 | 0.82 | 0.31 | 0.87 | 0.32 | 0.90 | 3.99 | 0.86 |
| W10114-3R | -1.14 | 0.82 | 0.35 | 0.87 | 0.33 | 0.90 | 2.48 | 0.86 |
| W10209-2R | -3.46 | 0.82 | 0.29 | 0.87 | 0.29 | 0.90 | 3.20 | 0.86 |
| W10209-7R | -0.41 | 0.82 | 0.32 | 0.87 | 0.32 | 0.90 | 17.05 | 0.86 |
| W6511-1R | -4.00 | 0.82 | 0.29 | 0.87 | 0.28 | 0.90 | 3.07 | 0.86 |
| W8886-3R | -3.01 | 0.68 | 0.32 | 0.77 | 0.31 | 0.84 | 4.19 | 0.76 |
| W8890-1R | -3.22 | 0.82 | 0.31 | 0.87 | 0.29 | 0.90 | 2.22 | 0.86 |
| W8893-1R | -4.70 | 0.82 | 0.32 | 0.87 | 0.29 | 0.90 | 3.24 | 0.86 |

**Table S3.** Variance components for clone (*G*), harvest environment (*E*, 109 vs. 138 DAP), and storage time (*T*, 0 vs. 6 weeks) in the 2016 experiment.

| Trait | Variance components |  |  |  |
| --- | --- | --- | --- | --- |
| | $V_G$ | $V_{GT}$ | $V_{GE}$ | $V_{GET}$ |
| Hue | 5.50 | 0.00 | 0.91 | 0.00 |
| Chroma | 5.50 | 0.00 | 0.91 | 0.00 |
| Lightness | 14.95 | 0.92 | 0.70 | 0.00 |

**Table S4.** Genotype BLUPs for each combination of harvest and storage time in the 2016 experiment.

| Clone<br>Storage time | Hue (°) |  |  |  | Chroma |  |  |  | Lightness |  |  |  |
| --- | --- | --- | --- | --- | --- | --- | --- | --- | --- | --- | --- | --- |
|  | At Harvest |  | 6 weeks |  | At Harvest |  | 6 weeks |  | At Harvest |  | 6 weeks |  |
|  | 109<br>DAP | 138<br>DAP | 109<br>DAP | 138<br>DAP | 109<br>DAP | 138<br>DAP | 109<br>DAP | 138<br>DAP | 109<br>DAP | 138<br>DAP | 109<br>DAP | 138<br>DAP |
| Chieftain | 1.64 | 3.39 | 5.09 | 5.96 | 0.33 | 0.35 | 0.31 | 0.33 | 0.36 | 0.38 | 0.32 | 0.36 |
| DarkRedNorland | -5.14 | -5.38 | -1.92 | -2.47 | 0.34 | 0.36 | 0.31 | 0.33 | 0.34 | 0.36 | 0.30 | 0.34 |
| RedEndeavor | -1.61 | -1.80 | 2.18 | 1.53 | 0.32 | 0.34 | 0.30 | 0.31 | 0.33 | 0.36 | 0.30 | 0.35 |
| RedLaSoda10 | 0.62 | 0.59 | 4.26 | 4.15 | 0.33 | 0.35 | 0.32 | 0.34 | 0.34 | 0.38 | 0.31 | 0.35 |
| RedNorland | -1.16 | -0.11 | 2.59 | 2.56 | 0.35 | 0.35 | 0.33 | 0.33 | 0.39 | 0.40 | 0.34 | 0.37 |
| RedPrairie | -1.43 | -0.74 | 1.82 | 2.53 | 0.35 | 0.37 | 0.33 | 0.34 | 0.37 | 0.40 | 0.32 | 0.37 |
| VillettaRose | -4.19 | -3.05 | -0.63 | 0.12 | 0.31 | 0.32 | 0.29 | 0.29 | 0.32 | 0.34 | 0.30 | 0.33 |
| W10114-3R | -3.70 | -2.59 | -0.13 | 0.09 | 0.35 | 0.37 | 0.31 | 0.33 | 0.33 | 0.36 | 0.30 | 0.34 |
| W10209-2R | -3.82 | -4.61 | -0.35 | -1.56 | 0.29 | 0.31 | 0.26 | 0.28 | 0.29 | 0.31 | 0.27 | 0.30 |
| W10209-7R | -2.04 | -0.55 | 1.46 | 2.05 | 0.30 | 0.33 | 0.28 | 0.30 | 0.30 | 0.33 | 0.27 | 0.31 |
| W6511-1R | -3.20 | -4.81 | 0.78 | -2.03 | 0.27 | 0.29 | 0.24 | 0.26 | 0.28 | 0.30 | 0.26 | 0.28 |
| W8890-1R | -4.32 | -4.61 | -1.04 | -1.24 | 0.30 | 0.33 | 0.28 | 0.30 | 0.29 | 0.31 | 0.27 | 0.30 |
| W8893-1R | -5.32 | -6.70 | -1.82 | -3.92 | 0.31 | 0.34 | 0.27 | 0.31 | 0.29 | 0.31 | 0.27 | 0.30 |

## REFERENCES

Andersen, A.W., C. Tong and D.E. Krueger. 2002. Comparison of periderm color and anthocyanins of four red potato varieties. American Journal of Potato Research 79: 249–253.

Bussan, A.J., V.M. Colquhoun, A.J. Davis, R.L. Gevens, D.J. Groves, B.M. Heider, et al. 2015. Commercial Vegetable Porduction in Wisconsin.

Butler, D., B. Cullis, A. Gilmour and B. Gogel. 2009. ASReml-R reference manual Version 3, Queensland Department of Primary Industries and Fisheries, Brisbane.

Clark, S.A., J.M. Hickey, H.D. Daetwyler and J.H.J. Van der Werf. 2012. The importance of information on relatives for the prediction of genomic breeding values and the implications for the makeup of reference data sets in livestock breeding schemes. Genet Sel Evol 44: 1–9.

Darrigues, A., J. Hall, E. Van Der Knaap and D.M. Francis. 2008. Tomato Analyzer-color Test: A New Tool for Efficient Digital Phenotyping. Journal of American Society of Horticulture 133: 579–586.

Gao, Y., J. Geng, X. Rao and Y. Ying. 2016. CCD-Based skinning injury recognition on potato tubers (*Solanum tuberosum L.*): A comparison between visible and biospeckle imaging. Sensors 16.

Halderson, J.L. and R.C. Henning. 1993. Measurements for determining potato tuber maturity. American Potato Journal 70: 131–141.

Henderson, C.R. 1975. Best Linear Unbiased Estimation and Prediction under a Selection Model. Biometrics 31: 423–447.

Hung, C., J. Murray, S. Ohmann and C. Tong. 1997. Anthocyanin accumulation during potato tuber development. Journal of American Society of Horticulture 122: 20–23.

Johansen, R.H., N. Sandar, W.G. Hoyman and E.P. Lana. 1959. Norland a new red-skinned potato variety with early maturity and moderate resistance to common scab. American Potato Journal 36: 12–15.

Jung, C., H. Griffiths, D. De Jong, S. Cheng, M. Bodis, T. Kim, et al. 2009. The potato developer (D) locus encodes an R2R3 MYB transcription factor that regulates expression of multiple anthocyanin structural genes in tuber skin. Theoretical and Applied Genetics 120: 45–57. doi:10.1007/s00122-009-1158-3.

Li, B., X. Xu, J. Han, L. Zhang, C. Bian, L. Jin, et al. 2019. The estimation of crop emergence in potatoes by UAV RGB imagery. Plant Methods 15: 15.

Lulai, E.C. 2007. The canon of potato science: 43. Skin-set and wound-healing/suberization. Potato Research 50: 387–390.

Lulai, E.C. and T.P. Freeman. 2001. The Importance of Phellogen Cells and their Structural Characteristics in Susceptibility and Resistance to Excoriation in Immature and Mature Potato Tuber (Solanum tuberosum L.) Periderm. Analysis of Botany 88: 555–561.

Lulai, E.C. and P.H. Orr. 1993. Determining the Feasibility of Measuring Genotyping Differences in Skin-Set. American Journal of Potato Research 70: 599–609.

Miller, N.D., N. Haase, J. Lee, S. Kaeppler, N. De Leon and E. Spalding. 2017. A robust, high-throughput method for computing maize ear,cob, and kernel attributes automatically from images. The Plant Journal 89: 169–178.

Moore, C., L. Johnson, I.-Y. Kwak, M. Livny, K. Broman and E. Spalding. 2013. High-Throughput Computer Vision Introduces the Time Axis to a Quantitative Trait Map of a Plant Growth Response. Genetics 195: 1077–1086.

National Potato Council. 2018. Potato Statistical Yearbook 2018.

Neubauer, J., E. Lulai, A. Thompson, J. Suttle, M. Bolton and L. Campbell. 2013. Molecular and cytological aspects of native periderm maturation in potato tubers. Journal of Plant Physiology 170: 413–423.

Patel, K.K., A. Kar, S.N. Jha and M.A. Khan. 2012. Machine vision system: a tool for quality inspection of food and agricultural products. Journal of Food and Science Technology 49: 123–141.

R. Core Team. 2018. R: A language and environment for statistical computing. Viena, Austria.

Reeve, R.M., E. Hautala and M.L. Weaver. 1969. Anatomy and compositional variation whitin potatoes. American Journal of Potato Research 46: 361–373.

Roe, M., J. Carlson, T. McManimon, A. Hegeman and C. Tong. 2014. Differential Accumulation and Degradation Of Anthocyanins In Red Norland Periderm is Dependent On Soil Type And Tuber Storage Duration. American Journal of Potato Research 91: 696–705. doi:10.1007/s12230-014-9402-z.

Rosen, C., J.A. Roessler, P.D. Petracek, C.B.S. Engelman and C. Tong. 2009. 2,4-Dichlorophenoxyacetic acid increases peonidin derivates in Red Norland periderm. American Journal of Potato Research 86: 15–23.

Schneider, C., W. Rasband and K. Eliceiri. 2012. NIH Image to ImageJ: 25 years of image analysis. Nature Methods 9.

Smith, A. 1978. Color gamut transform pairs. Computer Graphics 12: 12–19.

Sulc, M., Z. Kotikova, L. Paznocht, V. Pivec, K. Hamouz and J. Lachman. 2017. Changes in anthocyanidin levels during the maturation of color-fleshed potato (*Solanum tuberosum L*.) tubers. Food Chemistry 237: 981–988.

Tao, Y., P. Heinemann, Z. Varghese, C. Morrow and H. Soomer III. 1995. Machine vision for color inspection of potatoes and apples. American Society of Agricultural Engineers 38: 1555–1561.

USDA, U.S.D.o.A. 2008. United States Standards for Grades of Potatoes. In: A. M. Service, editor.

Vander Haeghen, Y. 2007. Chart_White_Balance, C4Real. Realistic, web-ready sRGB pictures from your digital camera.

Vulavala, V., E. Fogelman, L. Rozental, A. Faigenboim, Z. Tanami, O. Shoseyoc, et al. 2017. Indetification of genes related to skin development in potato. Plant Molecular Biology 94: 481–494.

Waterer, D. 2010. Influence of growth regulators on skin colour and scab diseases of red-skinned potatoes. Canadian Journal of Plant Science 90: 745–753.

Zhang, Y., C.S. Jung and W.S. De Jong. 2009. Genetic analysis of pigmented tuber flesh in potato. Theoretical and Applied Genetics 119: 143.

